# The use of disinfectant in barn cleaning alters microbial composition and increases carriage of *Campylobacter jejuni* in broiler chickens

**DOI:** 10.1101/2021.10.29.466552

**Authors:** Yi Fan, Andrew J. Forgie, Tingting Ju, Camila Marcolla, Tom Inglis, Lynn M. McMullen, Benjamin P. Willing, Douglas R. Korver

## Abstract

To maintain food safety and flock health in broiler chicken production, biosecurity approaches to keep chicken barns free of pathogens are important. Canadian broiler chicken producers must deep clean their barns with chemical disinfectants at least once annually (full disinfection; FD) and may wash with water (water-wash; WW) throughout the year. However, many producers use FD after each flock, assuming a greater efficacy of more stringent cleaning protocols, although little information is known regarding how these two cleaning practices affect pathogen population and gut microbiota. In the current study, a cross-over experiment over four production cycles was conducted in seven commercial chicken barns to compare WW and FD. We evaluated the effects of barn cleaning method on the commercial broiler performance, cecal microbiota composition, pathogen occurrence and abundance, as well as short-chain fatty acid concentrations in the month-old broiler gut. The 30-day body weight and mortality rate were not affected by the barn cleaning methods. The WW resulted in a modest but significant effect on the structure of broiler cecal microbiota (weighted-UniFrac; adonis *p* = 0.05, and unweighted-UniFrac; adonis *p* = 0.01), with notable reductions in *Campylobacter jejuni* occurrence and abundance. In addition, the WW group had increased cecal acetate, butyrate and total short-chain fatty acid concentrations, which were negatively correlated with *C. jejuni* abundance. Our results support the use of WW over FD to enhance the activity of the gut microbiota and potentially reduce zoonotic transmission of *C. jejuni* in broiler production without previous disease challenges.

## Introduction

In broiler chicken production, biosecurity measures are important to maintain flock health and food safety. Regulations of the current on-farm-food safety assurance program require Canadian broiler chicken producers to clean barns with disinfectants at least annually (1). Barn cleanouts within the year can be done without disinfectant using a water wash (**WW**); however, in practice many producers continue to perform full disinfection (**FD**) using chemical disinfectants after each flock. Using chemical disinfectants removes a high proportion of microbes (2), and, may reduce the transmission of beneficial microbes between flocks that can outcompete pathogens in the environment and thus select for disinfectant-resistant pathogens that further increases the risk of pathogen contamination of animal products (3). To date, little information is available regarding how these cleaning measures affect chicken health and food safety. Furthermore, emerging evidence has shown the importance of commensal microbial community to nutrient metabolism, feed efficiency (4–6), host resistance to pathogens (7), and immune system development (8, 9). In chicken production, the establishment of a symbiotic microbiota has been shown to improve nutrient utilization (10), and prevent disease development (11). Therefore, establishing healthy host-microbe interactions early in production may provide a possible solution to help maintain, or even enhance broiler performance in an antimicrobial growth promoter-free environment. The gut microbiota assembly and development is significantly influenced by the initial environment to which chicks are exposed (12). Placing birds in a WW barn, thus allowing an early exposure to bacteria from a previous healthy flock, may advance the development of a commensal microbiota in the broiler chickens.

Previous work has shown that exposure to reused litter altered early-life gut microbiota and increased infection resistance to pathogens in broiler chickens (13–15). Reused litter induced changes in the gut microbiota of chicks over the first two weeks of life, resulting in an increased predominance of *Clostridiales* in the gut (16). More recently, the use of recycled litter was reported to increase the predominance of some potentially beneficial bacteria, such as *Faecalibacterium*, a short-chain fatty acid (**SCFA**) producer whose increased abundance in young chick ceca continued as the chicken matured (17). Commensal microbes and SCFAs are important in maintaining gut homeostasis. For example, butyrate increases intestinal epithelial oxygen consumption, which helps to maintain an anaerobic environment (18). SCFAs also modulate host immune responses by suppressing pro-inflammatory cytokine expression to achieve homeostasis (19).

Both poultry and zoonotic pathogens have long been a major concern to the poultry industry. The poultry gut microbiota plays an important role in pathogen exclusion. For example, commensal members of the chicken gut microbiota inhibited *Salmonella enterica* colonization by competitive exclusion (15, 20). Colonizing young chicks with a diverse set of adult chicken commensal isolates reduced *Campylobacter jejuni* colonization (21). In addition to the potential impact on human health by zoonotic diseases, the economic impact of poultry diseases is a major concern of chicken producers. Bacterial pathogens such as *Clostridium perfringens* can cause necrotic enteritis, resulting in reduced growth and feed efficiency, and in severe cases, increased mortality (22). Research has demonstrated that using a cocktail of probiotics reduced the level of cecal *C. perfringens* and associated intestinal lesions (23). However, antibiotics have been the most widely used approach to keep the prevalence of *C. perfringens* infections low in industry (24). Due to legislation and consumer pressure to reduce antibiotic use in animal production, it is important to find effective alternatives to control necrotic enteritis.

To date, limited information is available on how WW and FD affect pathogen prevalence, gut microbial communities, nutrient metabolism, and host responses in chickens. In this study, a cross-over experiment was designed using seven similar commercial chicken barns to compare WW and FD over the course of four production cycles at two locations. We evaluated the effects of barn cleaning method on the commercial broiler intestinal microbiota, occurrence of select pathogens and abundance as well as SCFA profile in 30-day-old broiler ceca.

## Materials and methods

### Broiler production house and barn cleaning

A commercial broiler company in Alberta, Canada provided all the chickens and facilities including a total of seven similarly engineered single-storey, cement-floored production houses at two locations for this study. The broiler facilities were environmentally-controlled metal houses with solid sidewall ventilation. Four alternating water and feed lines ran the entire length of each house.

For FD, chicken manure, used litter, and organic matter were completely removed from the chicken house after depopulation followed by a two-step disinfection: 1) All surfaces were thoroughly covered with foam containing 7% sodium hydroxide, 7% 2-(2-2-butoxyethoxy) ethanol, 6% sodium laureth sulfate, 5% sodium N-lauroyl sarcosinate, and 5% tetrasodium ethylenediaminetetraacetic acid for 60 mins, and subsequently rinsed with high-pressure followed by low-pressure water-wash with the water temperature set at 35℃; 2) after the broiler house was air dried, all surfaces were covered with foam containing 10% glutaraldelhyde, 10% benzalkonium chloride, and 5% formic acid for 60 mins followed by high-pressure water rinse. After the two-step disinfection, broiler houses were left to air-dry overnight followed by placement of fresh wood shavings. For WW, manure and used litter were removed, followed by low-pressure water rinse with the water temperature set at 35℃ of the facility surfaces, air dried, and placement of fresh wood shavings. The current study was performed on 28 production flocks, and the FD and WW treatments were each applied on 14 production flocks. For each chicken barn, two flocks of each treatment were assigned according to the schedule shown in Table 1.

**Table 1.** Production cycles, barns and cleaning treatment schedule.

|  | Barn A | Barn B | Barn C | Barn D | Barn E | Barn F | Barn G |
| --- | --- | --- | --- | --- | --- | --- | --- |
| Cycle 1 | FD | FD | FD | FD | FD | WW | WW |
| Cycle 2 | FD | FD | WW | WW | FD | WW | WW |
| Cycle 3 | WW | WW | WW | WW | WW | FD | FD |
| Cycle 4 | WW | WW | FD | FD | WW | FD | FD |

WW, water-wash; FD, full disinfection

### Chicken management and sample collection

Animal use for this experiment was approved by the Animal Care and Use Committee: Livestock of the University of Alberta following the Canadian Council on Animal Care guidelines (25). For each flock, 14,000 Ross 308 broiler chicks were placed at 1 day of age and confined to half of the house, then allowed access to the entire house at 7 days of age. All chickens were fed *ad libitum* and reared from 1 day of age through processing at about 32-35 days of age when the average target live weight of 4 pounds is reached. Each flock had a placement based on a maximum stocking density of 30 kg/m^2^. Overall mortality rate and body weight of birds at day 32 were recorded for each barn.

At day 30, five birds from each flock were euthanized using cervical dislocation for sampling. To ensure representative sampling, the five birds were each randomly selected from different areas within each barn.

Sample collections were conducted aseptically. Briefly, the sampling table was cleaned with 70% ethanol before and between every bird dissection. Tools and collection tubes were autoclaved and tubes were sealed until samples were added. Approximately 300 mg of cecal contents were collected and placed in sterile 2 ml Eppendorf tubes containing 1 ml of 0.1% buffered peptone water (Oxoid, Basingstoke, Hampshire, UK) for detection of cecal *Campylobacter* and *Salmonella* by enrichment. In addition, approximately 500 mg of cecal contents were snap frozen on dry ice and stored at −80 ℃ for subsequent DNA extraction.

### Cecal *Campylobacter* and *Salmonella* enrichment

For genus *Campylobacter* enrichment and detection, cecal contents were homogenized with sterile peptone water. Homogenized cecal contents were incubated at 37°C overnight followed by inoculation in Bolton *Campylobacter* selective broth (Oxoid, Basingstoke, Hampshire, UK) at 42°C for 24 h under microaerobic conditions (5% O_2_, 10% CO_2_, 85% N_2_). Aliquots (100 μl) were serially diluted and spread onto Preston *Campylobacter* selective agar (Oxoid, Basingstoke, Hampshire, UK). For *Salmonella* enrichment, cecal contents were homogenized with universal pre-enrichment broth (Sigma-Aldrich, Oakville, ON, Canada) and incubated aerobically at 35°C for 24 h. The cultured broth was transferred to 10 ml of tetrathionate broth and to 10 ml of selenite cystine broth. The tetrathionate broth and selenite cystine broth were incubated for 24 h at 42°C and 35°C, respectively. After incubation, tetrathionate and selenite cystine broths were streaked onto xylose-lysine-tergitol 4 agar (**XLT-4**; Difco, Becton, Dickinson and Company Sparks, MD, USA) and brilliant green sulfa agar (**BGS**; Difco, Becton, Dickinson and Company Sparks, MD, USA) plates, respectively. Plates of XLT-4 and BGS agar were incubated at 35°C for 24 h.

Occurrence score for detected pathogens was calculated to evaluate the effect of cleaning method on pathogen occurrence in the chicken ceca. Occurrence score was defined as the number of positive birds divided by the total number of birds sampled from the same flock within each barn.

### DNA extraction and microbiome analyses

Total DNA was extracted from cecal contents using the QIAamp Fast DNA Stool Mini Kit (Qiagen, Valencia, CA, USA) with an additional bead-beating step with ~200 mg of garnet beads at 6.0 m/s for 60 s (FastPrep-24 5G instrument; MP Biomedicals Inc., Santa Ana, CA, USA). Amplicon libraries were constructed according to the manufacture protocol from Illumina (16S Metagenomic Sequencing Library Preparation) targeting V3-V4 region of the 16S rRNA gene (primers: Forward: 5 ′ -TCGTCGG CAGCGTCAGATGTGTATAAGAGACAGCCTACGGGNGGCWGCAG-3 ′; Reverse: 5 ′ -GTCTCGTGGGCTCGGAGATGTGTATAAGAGACAGGACT ACHVGGGTATCTAATCC-3′). An Illumina MiSeq Platform (2×300 cycles; Illumina Inc., San Diego, CA, USA) was used for a paired-end sequencing run. All sequences were submitted to NCBI Sequence Read Archive under BioProject ID: PRJNA767330. The quality of sequencing reads were assessed using FastQC. Sequenced reads were processed using Quantitative Insight into Microbial Ecology2 (**QIIME2**) −2020.6 (26). Divisive Amplicon Denoising Algorithm 2 was used to denoise and generate paired-end representative read with truncation lengths of 280 bp forward and 260 bp reverse reads (27). Amplicon sequence variant (**ASV**) feature table was created based on the denoised results. Qiime2’s q2-feature-classifier was used to assign taxonomy (28) with a pretrained classifier “Greengenes 13_8” (99% identity) (29). Analyses of diversity were done by the ‘diversity core-metrics-phylogenetic’ command normalizing to a sampling depth set by the sample with the lowest number of reads (17,309). Chao1 and Shannon diversity indices were calculated with ‘diversity alpha-phylogenetic’. Significance of alpha diversity was determined by ‘diversity alpha-group-significance’. Beta diversity was determined in QIIME2 using the unweighted- and weighted-Unifrac distance metric and a principal coordinate analysis (**PCoA**) was plotted using phyloseq package in R (version 3.6.1). Permutational Multivariate Analysis of Variance (**PERMANOVA**) based on the unweighted- and weighted-UniFrac distance matric was used to determine whether there were significant differences in community structures between treatments (adonis function). Differentiate taxa relative abundance between treatments was determined by the analysis of composition of microbiome (**ANCOM**) in QIIME2 (26).

### Quantitative PCR (qPCR)

A qPCR assay was used to quantify *C. jejuni*, *C. perfringens* as well as total cecal bacteria using hippurate hydrolase (***HipO***), necrotic enteritis B-like toxin (***NetB***) and the targeted 16s rRNA gene, respectively (Table 2). Total genomic DNA was extracted from cecal contents as described above. The concentration of DNA was determined by a NanoDrop 2000c spectrophotometer (Thermo Fisher Scientific, Waltham, MA, USA). PerfeCTa SYBR Green Supermix (Quantabio, Beverly, MA, USA) was used for qPCR assays which were conducted on an ABI StepOne real-time System (Applied Biosystems, Foster City, CA, USA) following the setup of 95°C for 3 min and 40 cycles of 95°C for 10 s, 60°C for 30 s. To generate targeted gene standards, the 16s rRNA gene, *HipO* and *NetB* were amplified by PCR with primers as listed. Concentrations of the amplified gene fragments were determined by a Quant-iT™ PicoGreen™ dsDNA Assay Kit (Invitrogen, Waltham, MA, USA) and used for standard curve. Gene copy numbers were determined using the Relative Standard Curve Method, and normalized to the weight of cecal content used for bacterial DNA extraction.

**Table 2.**
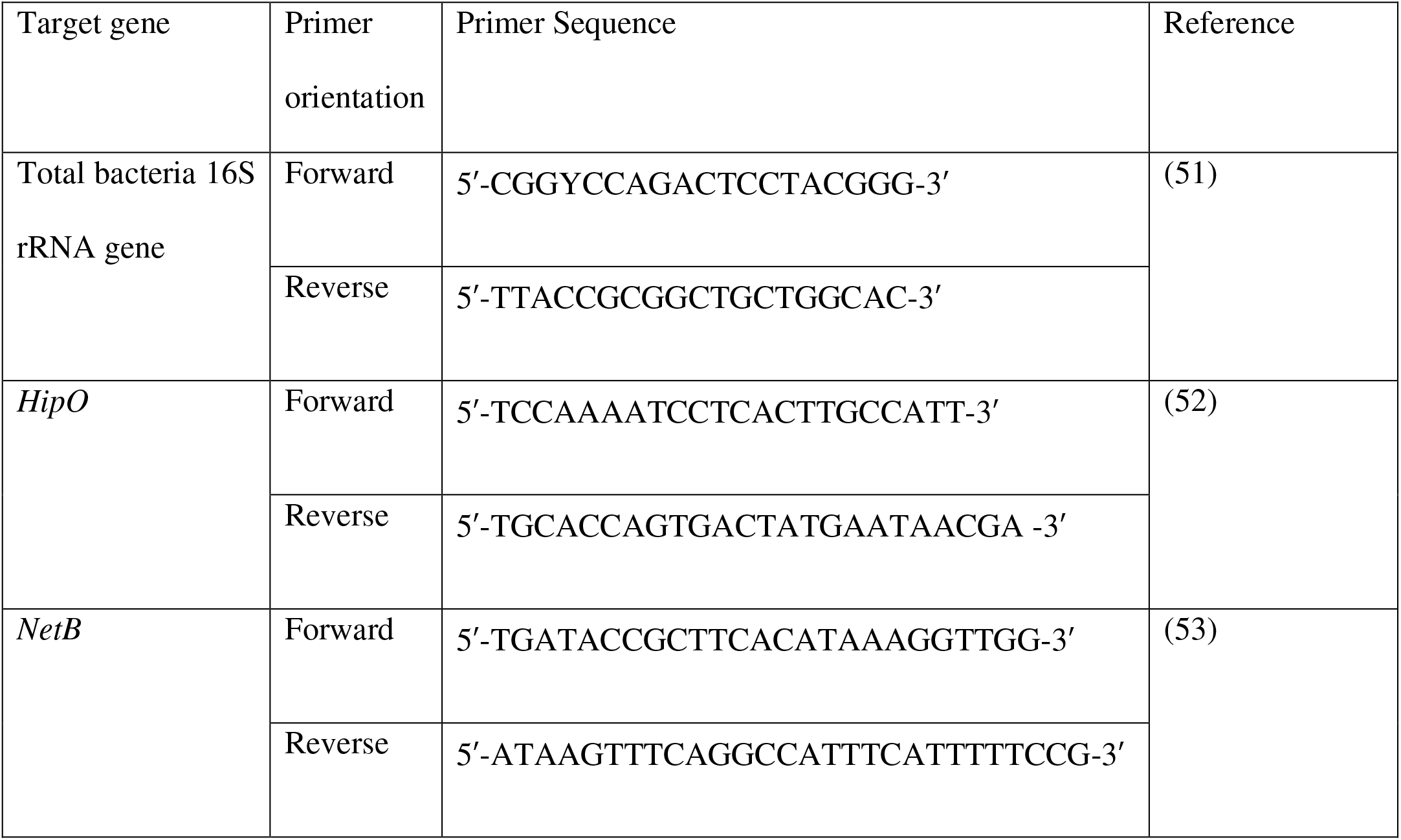
Primers used for qPCR assay of broiler chicken cecal samples collected at 30 days of age.

### Short chain fatty acid (SCFA) analysis

Approximately 30 mg per sample of snap frozen cecal content was weighed, followed by homogenization with 25% phosphoric acid. Samples were centrifuged at 21,130 x g for 10 min and the supernatant was collected and filtered using a 0.45 μm filter. Isocaproic acid (23 μmol/ml) was added at a 1:4 ratio to samples as an internal standard. Samples were analyzed on a Bruker Scion 456-Gas Chromatography instrument (Bruker, Billerica, MA, USA).

### Statistical analyses

Unless otherwise stated, statistical analyses were conducted using GraphPad Prism 8 (Graphpad Software, San Diego, CA, USA). Because no location effects were observed for any measurements (i.e. chicken performance and gut microbial structure), all data were analyzed based on treatment across location. Statistically significant differences were determined (*p* < 0.05) by an unpaired student’s t-test for parametric data (i.e., analyses of performance, qPCR and SCFAs). The Kruskal–Wallis test was used to determine the significance of non-parametric data (i.e., microbiome alpha diversity indices). The Spearman correlation was used to correlate SCFA concentration and bacterial relative abundance. Correlation significance was determined by psych package and visualized using corrplot package in R (version 3.6.1).

## Results

### Chicken 32-day body weight and mortality were not affected by the cleaning methods

The 32-day average body weight in the WW group was comparable to the FD group (Fig.1a, *p* = 0.22). In addition, no difference in 30-day mortality was observed between the two barn cleaning treatments (Fig1b, *p* = 0.91), suggesting that the cleaning method had a minimal impact on the flock performance.

**Figure 1.**
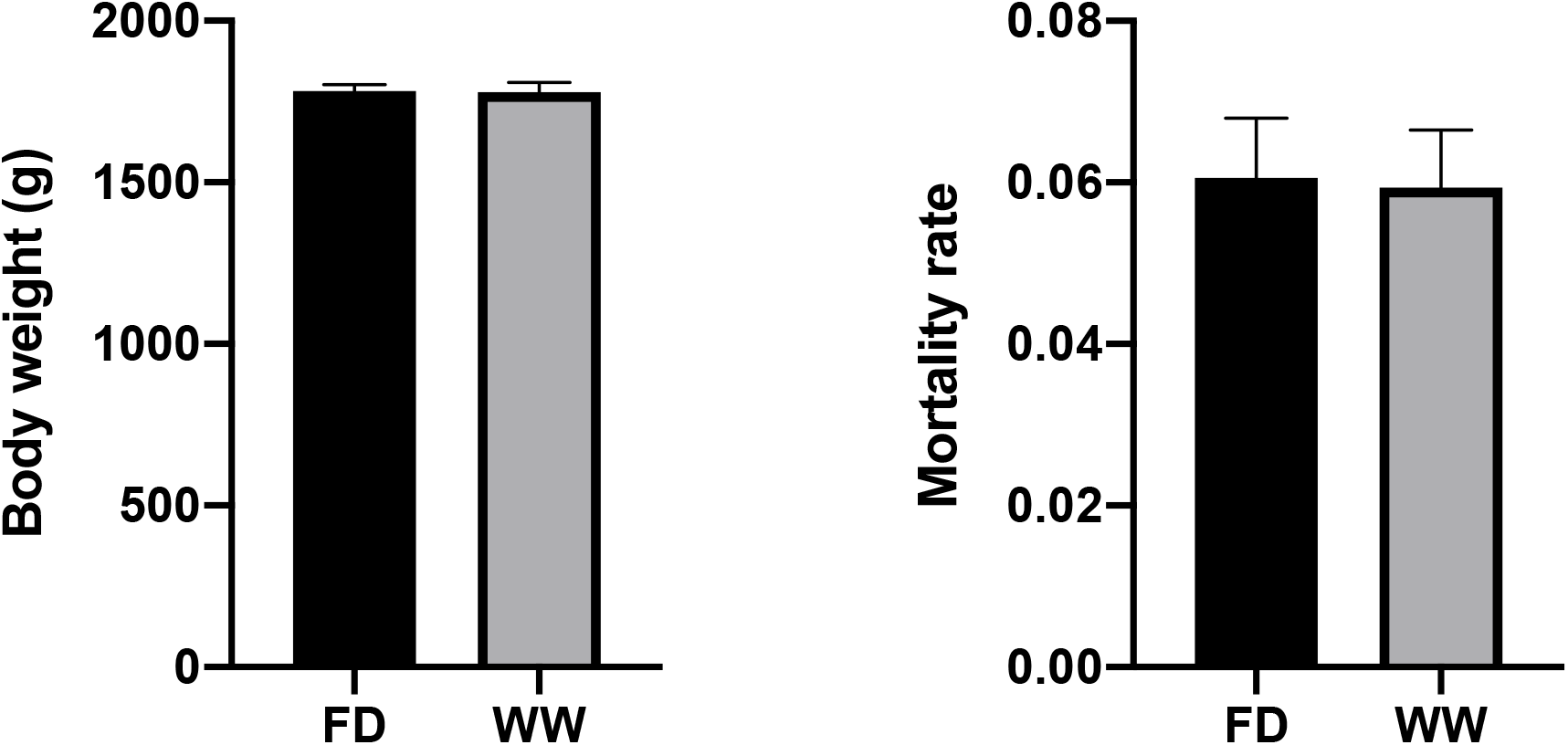
Broiler chicken flock performance at 32 d of age. (a) Flock mean body weight at day 30, 1782 ± 30.09 and 1780 ± 20.59 g for FD and WW, respectively (n=12 flocks/treatment, mean ± SEM; FD, full disinfection; WW, water-wash); (b) Flock mean mortality rate at day 30, 0.061 ± 0.0074 and 0.059 ± 0.0071 for FD and WW, respectively (n=14 flocks/treatment, mean ± SEM; FD, full disinfection; WW, water-wash).

### Chicken barn FD resulted in increased *Campylobacter* occurrence in the 30-day-old chicken ceca

*Salmonella* was not detected through enrichment in any of the samples collected through the study, therefore, the impact of treatment on *Salmonella* shedding could not be assessed. To evaluate the effect of cleaning method on the pathogen occurrence, an occurrence scoring method was used. In the *Campylobacter* enrichment assay, the WW group exhibited a significantly lower *Campylobacter* occurrence score compared to the FD group (*p* < 0.05) (Fig. 2). *Campylobacter* was detected in at least one sample from each FD flock. Therefore, the WW rearing environment reduced the occurrence of *Campylobacter* colonization in the 30-day-old chicken ceca.

**Figure 2.**
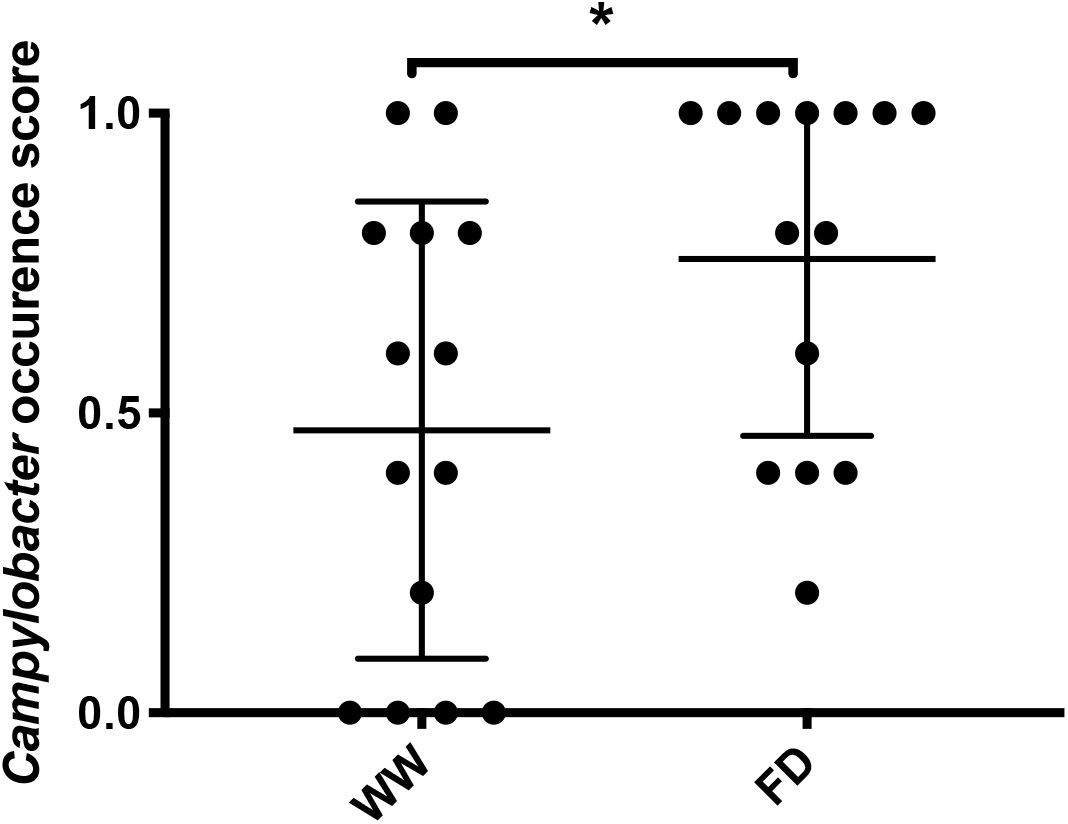
Broiler chicken cecal *Campylobacter* occurrence score at 30 d of age. Results showed the mean flock score ± SEM (n = 14/treatment, *, *p* < 0.05). *Campylobacter* occurrence score = number of pathogen positive birds/total number of birds sampled per barn. WW, water-wash; FD, full disinfection.

### Chicken barn FD increased *C. jejuni* abundance

The quantification of cecal microbial load using qPCR showed that the WW group did not differ from the FD group in cecal *C. perfringens* load (5.81 and 6.06 log10 copies/g of *netB* for WW and FD group, respectively, *p =* 0.20) (Fig. 3a). Consistent with the *Campylobacter* enrichment assay results, the WW group exhibited approximately 0.9-log10 lower *hipO* copy numbers compared to the FD group (Fig. 3b, 8.13 and 9.03 log10 copies/g of *hipO* for the WW and FD group, respectively, *p* < 0.05), indicating that the decreased sanitation stringency reduced *C. jejuni* colonization in the mature chicken ceca. Furthermore, the barn cleaning method did not affect the total bacterial load in the chicken ceca (Fig. 3c, *p* = 0.15).

**Figure 3.**
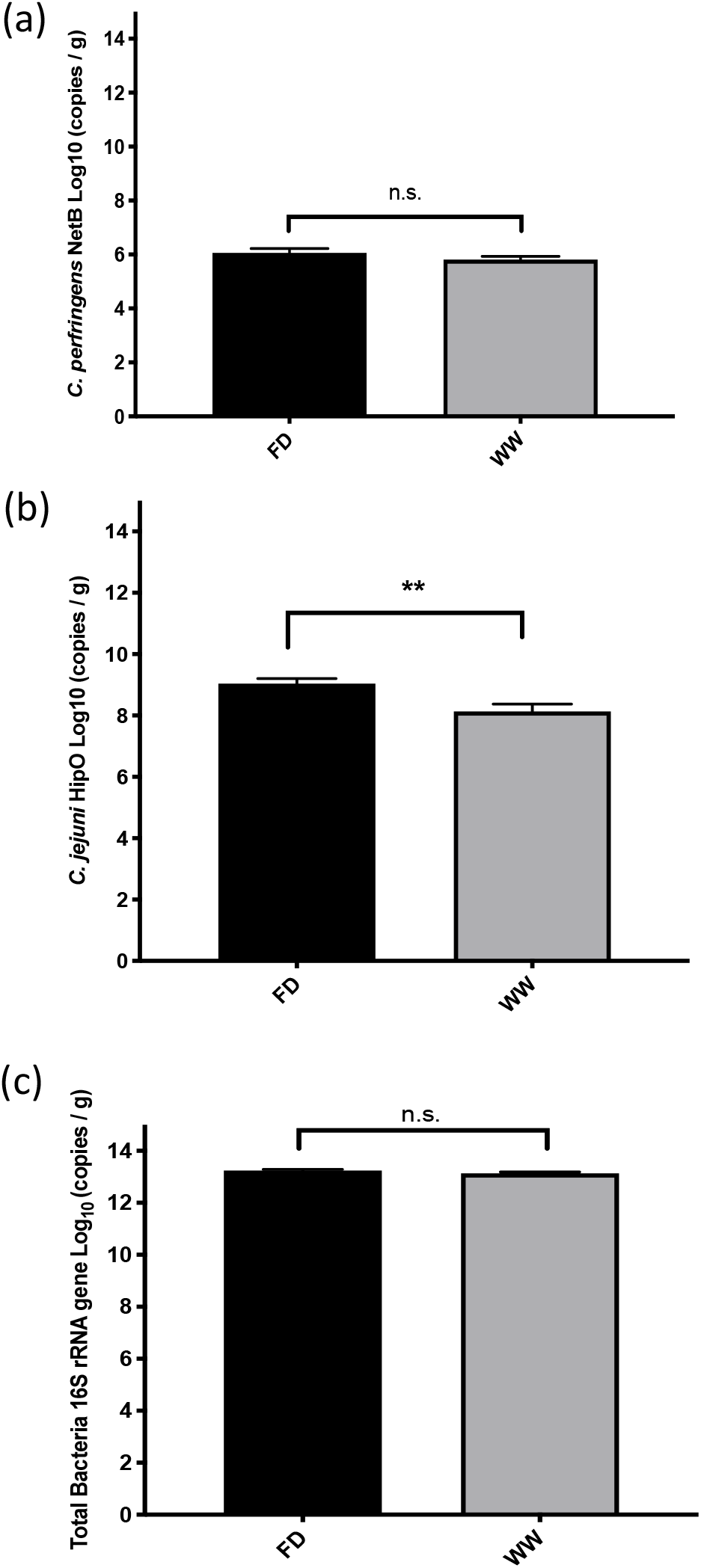
Broiler chicken (30 d of age) cecal qPCR targeting *C. perfringens* netB gene (a), *C. jejuni* hipO gene (b), and bacterial 16S rRNA gene (c). Results are the average copy number of each target gene (mean ± SEM, n=70/treatment, **, *p* < 0.01, n.s., *p* > 0.05). WW, water-wash; FD, full disinfection.

### Barn cleaning methods had subtle impacts on the 30-day-old chicken microbiome

On average, 42,943.3 ± 2757.8 (mean ± SEM) reads per sample were generated and processed by QIIME2 pipeline, resulting in a total of 3,845 ASVs. When focusing on the effect of different cleaning methods, the cecal microbiomes of broilers from the WW group and FD group had comparable richness and evenness indicated by alpha diversity indices (chao1, *p* = 0.71; and Shannon, *p* = 0.25) (Fig.4). Beta diversity analyses based on both weighted- and unweighted-Unifrac matrices suggested that cecal microbial communities in WW group differed from the FD group (adonis *p* = 0.05, R2 = 0.012 and adonis *p* < 0.01, R2 = 0.013 for weighted-Unifrac and unweighted-Unifrac matrix, respectively) (Fig. 5). Differences in abundances of two bacterial taxa were also suggested by ANCOM (Fig. 6). The genus *Helicobacter* was more predominant in the WW group (W = 85), whereas the family *Bacillaceae* was more predominant in the FD group (W = 66). These results suggested that the barn cleaning treatments influenced the relative abundance of two bacterial taxa, and in turn led to a modest but significant impact on overall structure of the chicken gut microbiota.

**Figure 4.**
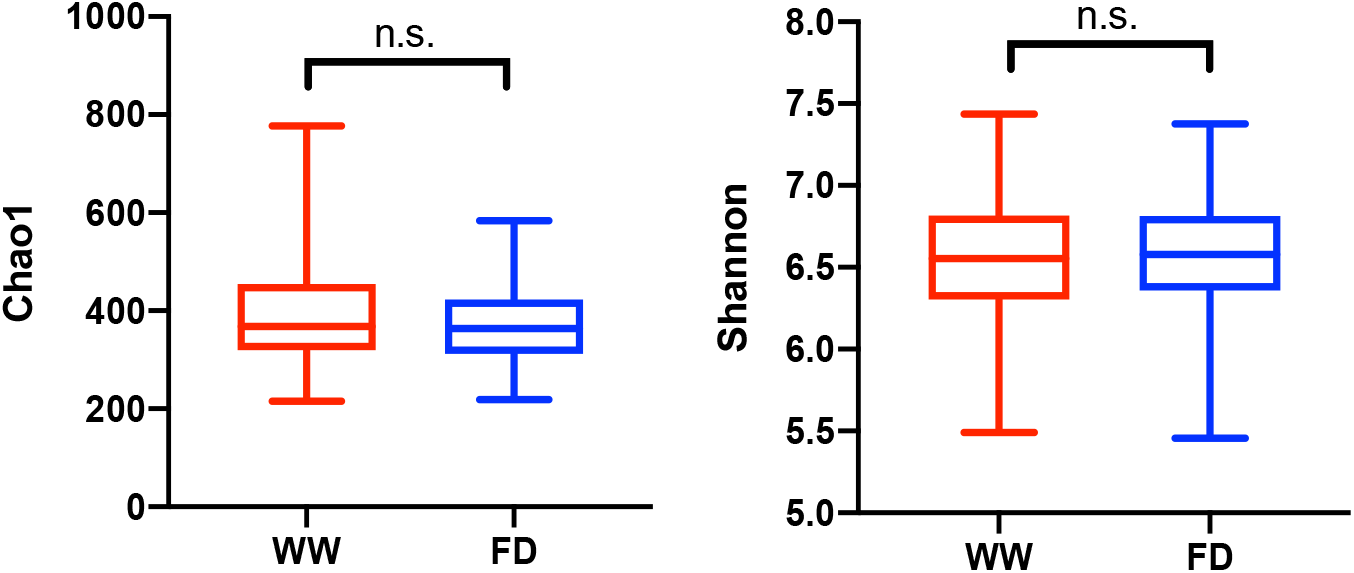
Alpha diversity of broiler chicken cecal microbiome at 30 d of age. Box-plots showing alpha diversity in samples using Chao1 index and Shannon index (n=70/treatment, n.s., *p* > 0.05). WW, water-wash; FD, full disinfection.

**Figure 5.**
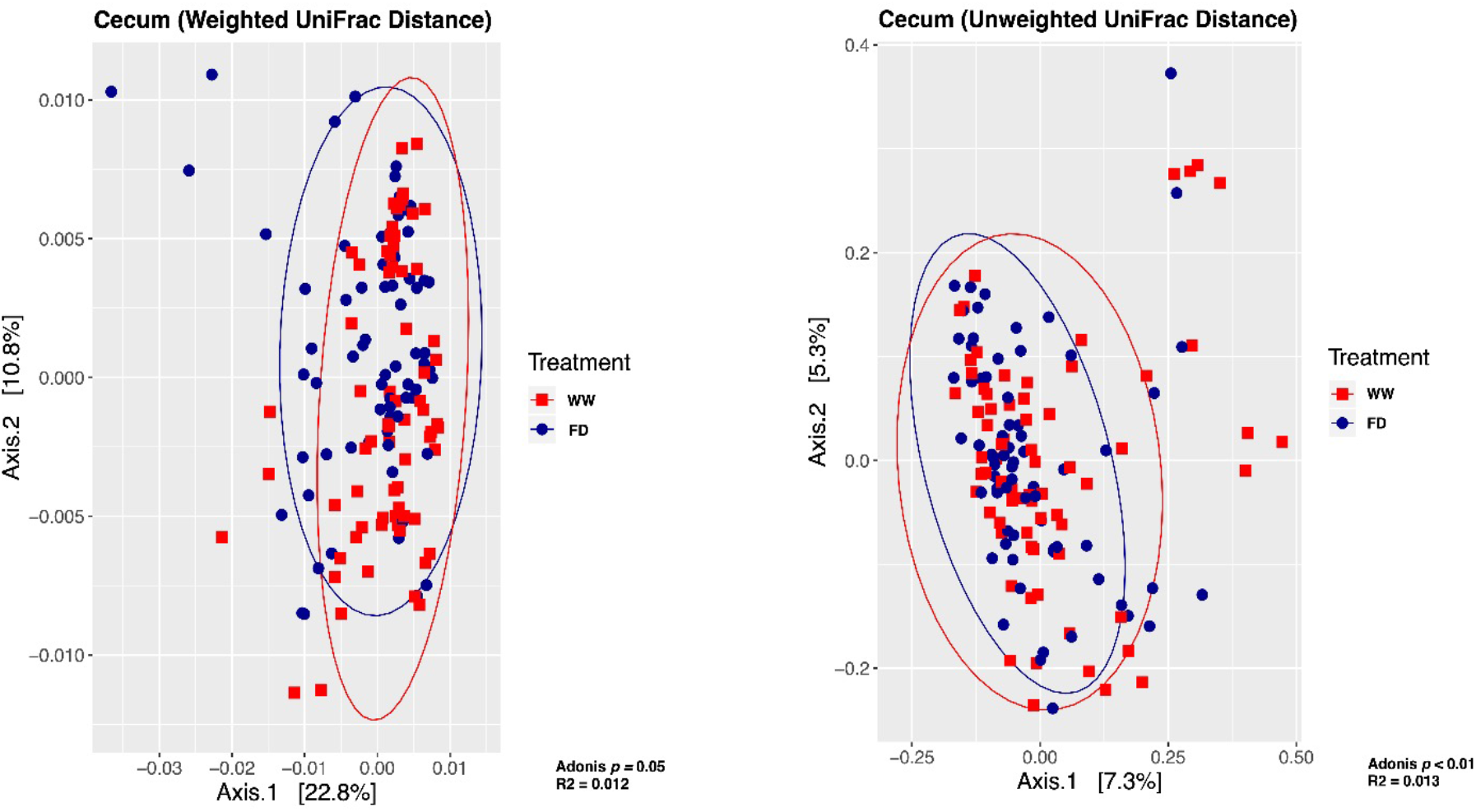
Principal coordinate analysis plots based on weighted- and unweighted-Unifrac distance matrices. Barn cleaning treatments had modest but significant effects on microbial community structure in the chicken ceca at 30 d of age (n=70/treatment). WW, water-wash; FD, full disinfection.

**Figure 6.**
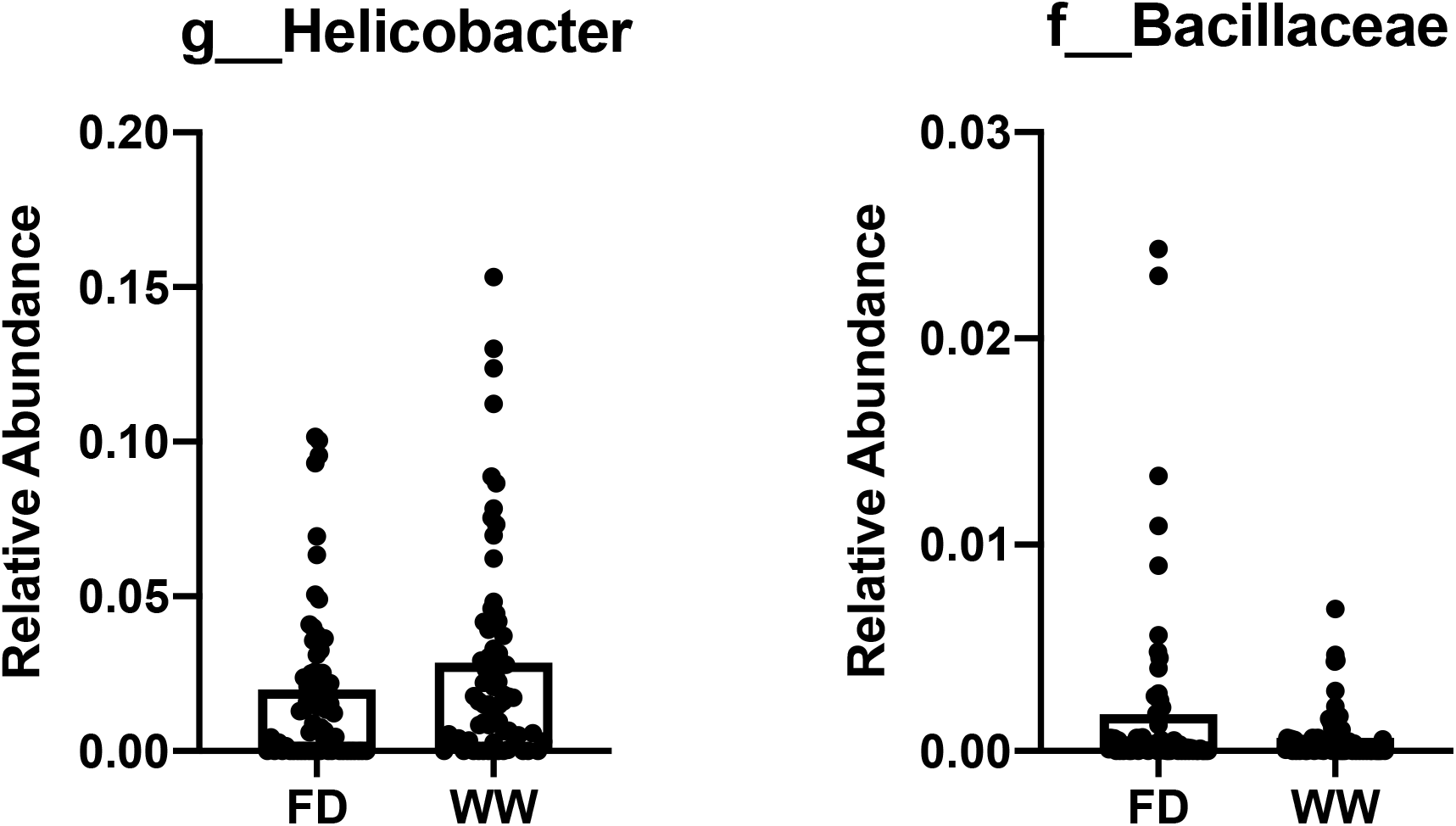
Relative abundance of genus *Helicobacter* and family *Bacillaceae.* The bar plot shows mean relative abundance of taxa of interests of each treatment with individual values (n=70/treatment). g, genus; f, family

### Cecal SCFA profile differed by cleaning methods

Cecal contents were subjected to gas chromatography to measure cecal SCFA concentration. The WW group showed significantly greater total SCFAs than the FD group (*p* < 0.01) (Fig. 7a). Specifically, acetate (Fig. 7b), propionate (Fig. 7c) and butyrate (Fig. 7d) concentration in the WW group were higher than that in the FD group. A trend for higher valerate concentration (Fig. 7e, *p=*0.06) was also observed in WW birds. Spearman correlation between SCFA concentration and bacterial relative abundance suggested a series of microbes that are correlated to the altered SCFA profile between treatments (Fig. 8). Total SCFAs and acetate concentration were negatively correlated with the genus *Campylobacter,* and members from orders RF32 and YS2 (*p* < 0.05). On the other hand, an unclassified genus belonging to the order of *Clostridiales* was positively associated to total SCFAs, acetate and butyrate concentrations (*p* < 0.05). Propionate concentration was negatively associated to the genus *Lachnospira* (*p* < 0.01) and an unclassified genus of the family *Enterobacteriaceae* (*p* < 0.05), whereas positively associated to the genus *Odoribacter* (*p* < 0.05), and an undetermined genus of the family *Clostridiaceae* (*p* < 0.05).

**Figure 7.**
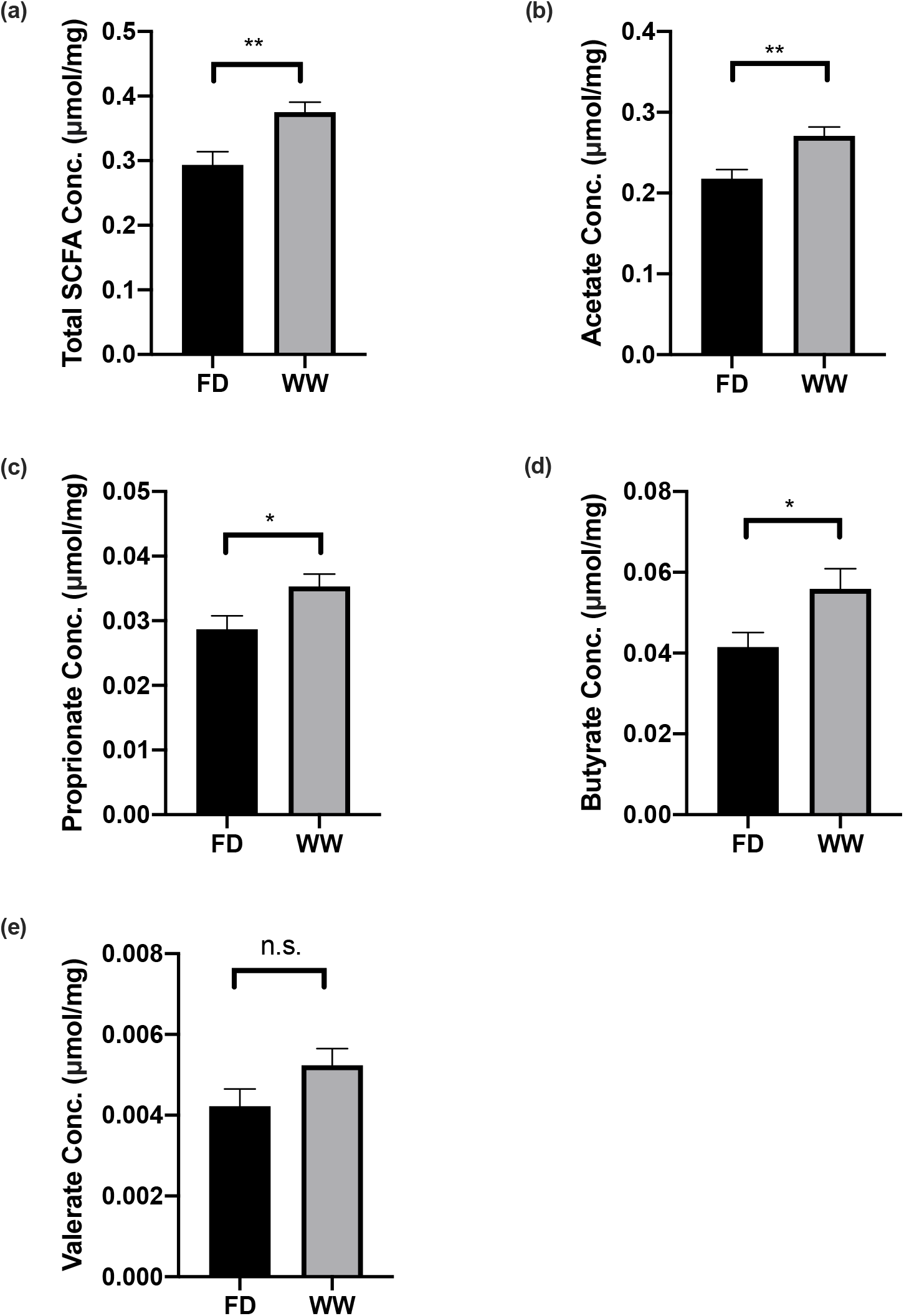
Cecal short chain fatty acid production in broiler chickens at 30 d of age. Results were shown as the average of (a) total SCFA concentration, (b) acetate, (c) propionate, (d) butyrate, and (e) valerate (mean ± SEM, n = 20/treatment, *, *p* <0.05, **, *p* < 0.01, n.s., *p* > 0.05). FD, full disinfection; WW, water-wash; Conc., Concentration.

**Figure 8.**
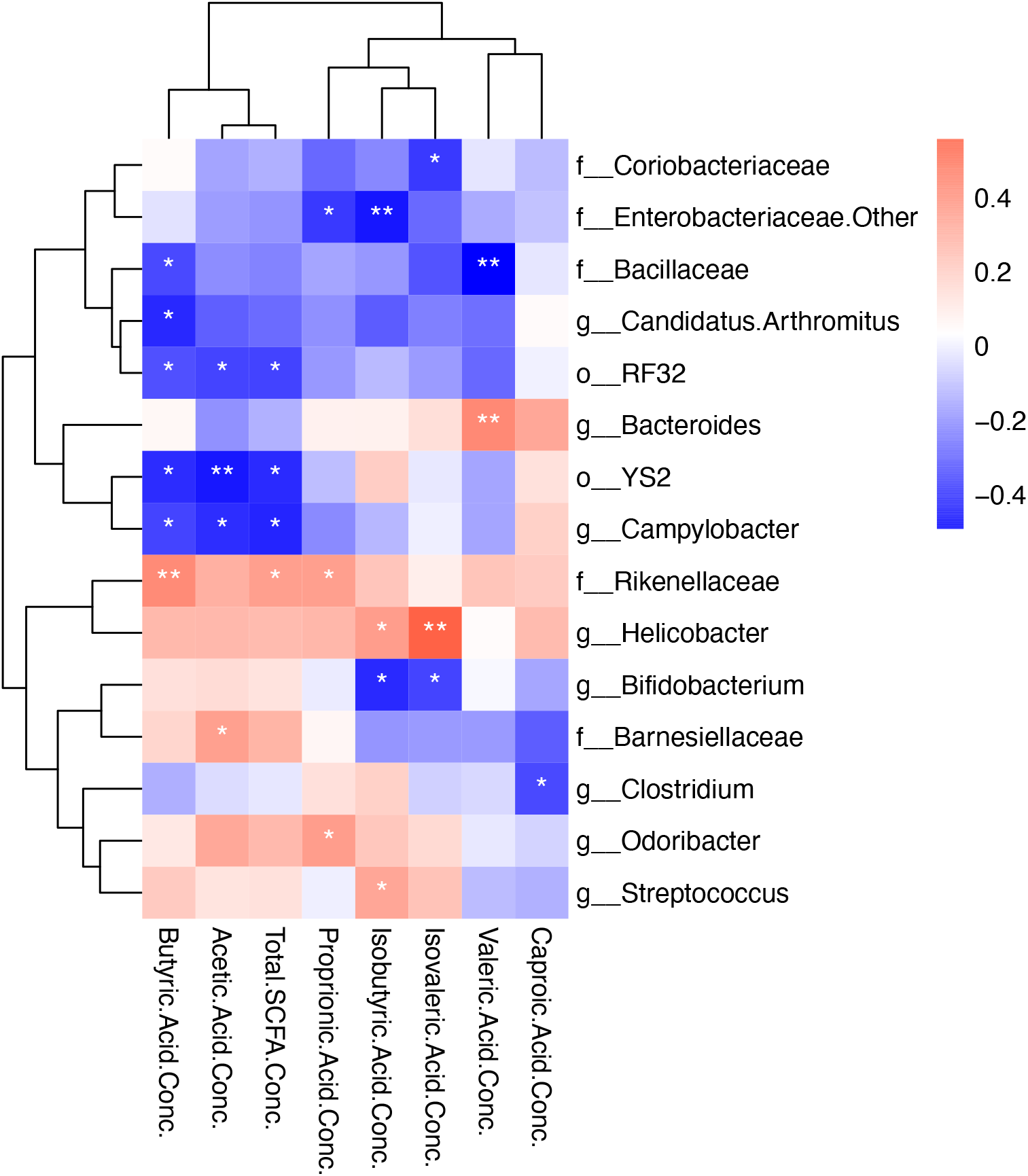
Heatmap showing Spearman correlations between cecal bacterial abundance and short chain fatty acid concentrations in broiler chickens at 30 d of age. *, *p* < 0.05; **, *p* < 0.01; Conc., Concentration.

## Discussion

This study was the first to characterize the impact of FD and WW on chicken gut microbiota in a commercial setting. In the absence of disease outbreak or pathogen challenge, we found that compared to FD, the WW did not have compromised flock performance. Furthermore, compared to FD, the WW of chicken barns has subtle but significant effects on the 30-day-old broiler gut microbiota and more importantly, resulted in decreased *C. jejuni* prevalence and abundance. This finding counters the intent of sanitation practices to reduce pathogen load and environmental transmission of zoonotic pathogens in poultry production.

Although limited research has explored how barn cleaning practices affect the development of chicken intestinal microbial communities, especially in the context of commercial production, efforts have been made to study the effect of reused litter on chicken gut microbiota. Some laboratory-scale research suggested that reused litter mainly influences broiler gut microbiota at early ages. Cressman et al. (2010) reported that compared to the used litter group, birds provided fresh litter had greater bacterial alpha diversity in ceca at 7 days of age (16). However, no treatment effects on the gut microbiota were observed in the later timepoints. In addition, chickens reared on fresh litter were colonized by microbes identified in fresh litter including *Lactobacillus*, unclassified *Lachnospiraceae* and *Enterococcus*, whereas chicks reared on used litter were colonized by typical poultry intestinal bacteria, such as members from the order *Clostridiales* (16). Similarly, Wang et al. (2016) reported that at both day 10 and day 35 of age, chicken gut microbiota was altered by the reused litter treatment with increased predominance of *Faecalibacterium prausnitzii* in ceca (17). In the current study, no treatment effect on alpha diversity of 30-day-old chickens was observed, and only a modest effect was shown on the beta diversity between FD and WW treatments. This may be explained by the more functional ecological niche provided by reused litter as compared with the WW group. Generally, reused chicken litter is a mixture of bedding material and excreta, which offers more surfaces and available nutrients for microbes to attach and survive on. In addition, as chickens are coprophagic, the consumption of litter material likely increases the opportunity for successful microbial transmission from one flock to the next (30).

*Helicobacter* was found to be less abundant in the FD group at day 30 compared to the WW group. Interestingly, *Helicobacter* is a genus identified as a disappearing member of the human gut microbiome, and may also be associated with increased use of disinfectants (31). In avian species, members in genus *Helicobacter* have been detected in wild birds (32). Studies on the relationship between *Helicobacter* spp. and chicken host are highly variable. Some members of genus *Helicobacter* are considered opportunistic pathogens in chickens. For example, *Helicobacter pullorum* infection was found to cause mild lesion in the chicken ceca (33). However, the effects of *Helicobacter* on host health can vary between different bacterial strains within species (34). Yin et al. (2018) reported that *Helicobacter* abundance increased in response to α-amylase, amylopectase and glucoamylase supplementation in a corn-based diet, and was associated with increased starch digestibility and higher mature bodyweight (35). In the current study, *Helicobacter* positively correlated to branched-chain fatty acids (**BCFA**s) isobutyrate and isovalerate (Fig. 7). BCFAs are often used as indicators of protein catabolism (36). Currently, the direct relationship between BCFAs and their impact on host health is still unclear (37). It is reported that BCFAs modulate adipocyte lipid and glucose metabolism, and contribute to increased insulin sensitivity (38). With 16S rRNA gene amplicon sequencing, it is difficult to discriminate bacteria to the species or strain level. Therefore, our identification of *Helicobacter* as the genus of increased predominance in the WW group needs to be further studied. In addition, information on metabolic functionality is also warranted to understand the role of *Helicobacter* in the chicken gut.

Interestingly the FD birds showed increases in the relative abundance of the family *Bacillaceae*. Members of the *Bacillaceae* family, such as *Bacillus* are Gram-positive, rod-shaped bacteria that can form endospores to survive in harsh physical and chemical environments (39). It is possible that the chemical conditions given by FD treatment provided a selective pressure that led to the increased level of *Bacillaceae*. Some members in this order have the ability to produce antimicrobial peptides, and are recognized as potential beneficial bacteria (40). Some *Bacillus* species, such as *Bacillus licheniformis* and *Bacillus subtilis*, have been commercially added into poultry feed as probiotics (41). However, not all *Bacillus* species are beneficial. *Bacillus cereus* is a food-borne pathogen that causes diarrhea (42). Recently, the prevalence of the *B. cereus* group made up 50% of *Bacillus* spp. isolates from retail chicken products (43). In addition, some non-*B. cereus* species were found to carry virulence genes and exhibited the same phenotypic virulence characteristics as *B. cereus*. Furthermore, tests of antimicrobial resistance have identified multi-drug resistant isolates regardless of virulence factors, indicating that further evaluation of the impact of *Bacillus* on food safety and public health is needed (43). In this sense, the effect of barn disinfection on increasing *Bacillaceae* may need to be carefully assessed.

The reduced *Campylobacter* load in the current study with WW treatment is consistent with previous studies showing that reused litter has the potential to reduce gut pathogen abundance in broiler chickens. It has been reported that reused litter reduced the colonization of *Salmonella enterica* serovar Typhimurium and *Salmonella enterica* serovar Enteritidis in infected birds (13, 44). In the current study, although *Salmonella* was not detected in any of samples by the enrichment assay, our results supported that the WW treatment did not increase *Salmonella* occurrence in the chicken intestine. On the other hand, the *Campylobacter* enrichment assay demonstrated that *Campylobacter* presented less frequently in the 30-day-old chicken ceca of the WW birds. Moreover, qPCR targeting *C.jejuni hipO* gene showed that the FD group had nearly one log more *hipO* gene copy numbers per gram of digesta compared to the WW group. The reduction of *C. jejuni* induced by the WW treatment associated with lower labor costs for cleaning (compared to FD) is a remarkable advantage from the production point of view. To date, limited data is available regarding the effects of cleaning treatments and disinfectants on *Campylobacter* occurrence and abundance in the chicken intestine. de Castro Burbarelli et al. (2017) examined the effects of poultry barn cleaning using neutral detergent versus a protocol using acidic and alkaline detergent with chemical disinfectants. Interestingly, there was a trend that *Campylobacter* was more frequently detected in the intestine of the stronger disinfection group (2), although the study was conducted in a controlled research setting. Furthermore, it has been shown that colonization of young chicks with bacterial cocktails of mature chicken commensal isolates reduced colonization by *C. jejuni* (45). Therefore, the FD treatment in the current study may have eliminated some microbes from the previous flock which can potentially compete with *Campylobacter*.

While the effects of cleaning method on the gut microbial composition were relatively modest, changes in concentrations of microbial metabolites, SCFAs, were observed. Total SCFAs, acetate and butyrate concentrations were higher in the WW group than their counterparts in the FD treatment. SCFAs enhance intestinal integrity as direct energy sources to enterocytes (46). Moreover, complex interactions between SCFAs, gut microbes, and the host immune system have been well documented (47). Briefly, fermented by the microbes from the lumen, SCFAs are imported into enterocytes and tissues via transporters and paracellular transport. SCFA receptors expressed on enterocytes and immune cells in the lamina propria and mucosal lymphoid tissue can activate signaling pathways to regulate host immune response according to the SCFA concentration to maintain intestinal homeostasis (47). Butyrate signaling through G-protein coupled receptors can confer anti-inflammatory properties in colonic dendritic cells by down-regulating the expression of cytokines and chemokines (19). In the present study, there were negative correlations between the relative abundance of *Campylobacter* and total SCFAs, acetate, and butyrate concentrations in the ceca. It has been suggested that SCFAs, especially butyrate, had shown bactericidal effect on *Campylobacter in vitro* (48). More recently, Awad et al. (2016) reported that *C.jejuni* infection led to reduced acetate and butyrate concentration in the chicken ceca (49). Adding microbeads coated with butyrate to feed was also found to reduce *Campylobacter* colonization in the chicken intestine (50). Together with our results, it is reasonable to suggest that the FD treatment discriminated against beneficial commensals in the gut environment, which could compete with or inhibit *Campylobacter* by producing SCFAs.

## Conclusion

We compared the effects of barn FD and WW methods on gut microbial community structures and pathogen prevalence of broiler chickens in a non-challenging commercial production setting. The results revealed that barn cleaning methods had little impact on the 30-day body weight and mortality rate of broiler chickens. In addition, the FD treatment had a subtle but significant effect on the broiler cecal microbiota with increased abundances of *C. jejuni* and decreased SCFA concentrations, which would support the adoption of WW as a standard practice. Thus, compared to FD, WW can be beneficial to broiler chicken production by inhibiting zoonotic pathogen colonization in the chicken gut with reduced cost and labor of cleaning. Further studies examining other barn disinfection practices and testing for other pathogens are warranted to identify the best practices to minimize pathogen load and maintain animal performance.

## Acknowledgements

This study was supported by the Alberta Livestock and Meat Agency, the Alberta Chicken Producers, and the Canadian Poultry Research Council. BPW is supported by the Canada Research Chair Program. Funders did not participate in study design, data collection and interpretation, or the decision to submit the work for publication.

We would also like to acknowledge Patrick Ward, Taresa Chieng, Tausha Prisnee, Koonphol Pongmanee, Steve Geerlinks, Johnathan Kielstra, Alejandro Rodriguez, Dr. Lukas Nickel, and Dr. Justina Zhang for contribution to the sample collection.

